# Single cell ATAC-seq in human pancreatic islets and deep learning upscaling of rare cells reveals cell-specific type 2 diabetes regulatory signatures

**DOI:** 10.1101/749283

**Authors:** Vivek Rai, Daniel X. Quang, Michael R. Erdos, Darren A. Cusanovich, Riza M. Daza, Narisu Narisu, Luli S. Zou, John P. Didion, Yuanfang Guan, Jay Shendure, Stephen C.J. Parker, Francis S. Collins

**Affiliations:** Department of Computational Medicine & Bioinformatics, University of Michigan, Ann Arbor MI 48109; National Human Genome Research Institute, National Institutes of Health, Bethesda MD 20892; Department of Genome Sciences, University of Washington, Seattle WA 98109; Department of Human Genetics, University of Michigan, Ann Arbor MI 48109; Department of Cellular and Molecular Medicine, University of Arizona, Tucson AZ 85721; Department of Biostatistics, Harvard School of Public Health, Boston MA 02115

**Keywords:** Islet, Epigenomics, Deep Learning, Single cell, Chromatin, Type 2 diabetes

## Abstract

**Objective:** Type 2 diabetes (T2D) is a complex disease characterized by pancreatic islet dysfunction, insulin resistance, and disruption of blood glucose levels. Genome wide association studies (GWAS) have identified >400 independent signals that encode genetic predisposition. More than 90% of the associated single nucleotide polymorphisms (SNPs) localize to non-coding regions and are enriched in chromatin-defined islet enhancer elements, indicating a strong transcriptional regulatory component to disease susceptibility. Pancreatic islets are a mixture of cell types that express distinct hormonal programs, and so each cell type may contribute differentially to the underlying regulatory processes that modulate T2D-associated transcriptional circuits. Existing chromatin profiling methods such as ATAC-seq and DNase-seq, applied to islets in bulk, produce aggregate profiles that mask important cellular and regulatory heterogeneity.

**Methods:** We present genome-wide single cell chromatin accessibility profiles in >1,600 cells derived from a human pancreatic islet sample using single-cell-combinatorial-indexing ATAC-seq (sci-ATAC-seq). We also developed a deep learning model based on the U-Net architecture to accurately predict open chromatin peak calls in rare cell populations.

**Results:** We show that sci-ATAC-seq profiles allow us to deconvolve alpha, beta, and delta cell populations and identify cell-type-specific regulatory signatures underlying T2D. Particularly, we find that T2D GWAS SNPs are significantly enriched in beta cell-specific and cross cell-type shared islet open chromatin, but not in alpha or delta cell-specific open chromatin. We also demonstrate, using less abundant delta cells, that deep-learning models can improve signal recovery and feature reconstruction of rarer cell populations. Finally, we use co-accessibility measures to nominate the cell-specific target genes at 104 non-coding T2D GWAS signals.

**Conclusions:** Collectively, we identify the islet cell type of action across genetic signals of T2D predisposition and provide higher-resolution mechanistic insights into genetically encoded risk pathways.

## 1. INTRODUCTION

Pancreatic islets consist of a cluster of at least five different endocrine cell-types (alpha, beta, delta, gamma, and epsilon), each producing a unique hormone in a distinct but coordinated manner [1]. Collectively, these clusters of cells work together to maintain insulin production and glucose homeostasis. Disruption of the complex interplay between the cell types, their organization, and their underlying regulatory interaction is known to be associated with type-2-diabetes (T2D) pathophysiology [2]. However, the exact cellular mechanisms through which different risk factors contribute to the disease risk are not completely understood. Using GWAS and eQTL mapping approaches, recent studies have discovered >400 independent signals (>240 loci) associated with T2D and T2D-associated traits [3], although remarkably, more than 90% of them localize to non-protein-coding regions of the genome [4]. Growing evidence suggests that many of these variants likely influence the RNA expression and cellular function of human pancreatic islets by altering transcription factor binding, critical components of a cellular regulatory network [5–9].

High-throughput epigenomic profiling methods such as ATAC-seq [10] and DNase-seq [11] have enabled profiling of chromatin accessibility across samples in a tissue-wide manner, providing the opportunity to identify millions of context-specific regulatory elements. However, these bulk-measurements of chromatin accessibility limit the precise understanding of how tissue heterogeneity and multiple cell-types in the population contribute to overall disease etiology [12]. Recent advances in single-cell transcriptomic and epigenomic profiling methods have enabled an unbiased identification of cell-type populations and regulatory elements in a heterogeneous biological sample. By mapping the chromatin-regulatory landscape at a single-cell resolution, recent single-nuclei studies have demonstrated the potential to discover complex cell populations, link regulatory elements to their target genes, and map regulatory dynamics during complex cellular differentiation processes [13–16]. The pancreatic islet gene expression landscape has been investigated at single-cell resolution in existing studies [17,18], but chromatin accessibility studies have been limited to fluorescence-activated cell sorting (FACS) methods for obtaining cell-type populations [19,20]. FACS based methods will miss identification of unknown or rarer cell-populations and are unable to produce pure cell-type populations due to reliance on the specificity of cell-surface markers [21,22].

Here, we present a genome-wide map of chromatin accessibility in >1,600 nuclei derived from a human pancreatic islet sample using single-nucleus-combinatorial-indexing ATAC-seq (sci-ATAC-seq) [23]. sci-ATAC-seq enables us to deconvolve cell populations and identify cell-type-specific regulatory signatures underlying T2D. Notably, we find that T2D GWAS SNPs are significantly enriched in beta cell-specific and cross cell-type shared islet open chromatin, but not in alpha or delta cell-specific open chromatin. We also demonstrate, using the less represented delta cell population (< 5% of total islet population), that deep learning can improve signal recovery and feature reconstruction for less abundant cell-populations using concepts borrowed from image upscaling methods. We anticipate that our deep learning method will enable analysis of heterogeneous tissues that may be harder to obtain in large numbers or contain rare sub-populations. Collectively, these results identify the islet cell-type of action across genetic signals of T2D predisposition and provide higher-resolution mechanistic insights into genetically encoded pathophysiology.

## 2. MATERIALS AND METHODS

### 2.1 Bulk Islet ATAC-seq

#### Sample processing

The human pancreatic islet samples were procured and processed as described by Varshney et al [8]. Briefly, the islets were obtained from the National Disease Research Interchange (NDRI) and processed according to the NHGRI institutional review board-approved protocols. The islet was shipped overnight from the distribution center. Upon receipt, we pre-warmed the islet to 37 degree in shipping media for 1-2h before harvest. ~50-100 islet equivalents (IEQs) were harvested and transposed in triplicate following the methods in Buenrostro et al [10]. The ATAC-seq library was barcoded and sequenced 2 × 125bp on a HiSeq 2000.

#### ATAC-seq analysis

Sequencing adapters were trimmed using cta (v0.1.2) [24] and aligned to hg19 reference genome using BWA-MEM (v0.7.15-r1140, options: -I 200,200,5000) [25]. Picard MarkDuplicates (v2.18.27) was used for duplicate removal and samtools [26] was used to filter for autosomal, properly-paired and mapped read pairs with mapping quality >= 30 (v1.9, options: -f 3 -F 3340 -q 30). Replicates across each sample were merged into a single file using samtools merge. For peak calling, each sample was downsampled to 25 million (M) reads and converted to BED file. We then used MACS2 [27] to call broad peaks (v2.1.1.20160309; options: --nomodel --broad --shift -100 -- extsize 200 --keep-dup all --SPMR) and removed those with FDR >0.05 and overlapping with ENCODE hg19 blacklists [28]. ATAC-seq coverage tracks were displayed using UCSC Genome Browser and Integrative Genomics Viewer (IGV). Summary statistics were calculated using Ataqv (v1.0) [29] and are available in interactive and downloadable format online (**Table S7**). For comparative purposes, we performed the same read trimming, alignment, filtering, downsampling, and peak calling steps on publicly available ATAC-seq data (**Table S1**). Peaks from each sample were merged to create a master peak set and Spearman correlation was computed on the RPKM normalized read-count matrix.

#### Determination of high-confidence peaks

We randomly sampled 2.5 M reads from each sample using samtools view and pooled them into one file so that each sample is equally represented. Peaks were called on the pooled file as discussed in the previous paragraph. We then determined the number of samples overlapping with each master peak using peaks called on individual samples.

#### Overlap of reads with chromHMM states

We tested for enrichment of ATAC-seq peaks across 13 islet-specific chromatin states using Genomic Association Tester (GAT) [30]. We ran GAT (v1.3.5, options: --number-samples 10,000) and filtered chromatin states with no significant enrichment (Bonferroni adjusted p-value < 0.05) of peaks in them. The log2 fold enrichment values across chromatin states were clustered using hierarchical clustering of the correlation matrix.

### 2.2. sci-ATAC-seq analysis

#### Sample processing

We used the combinatorial cellular indexing method to generate single-nuclei chromatin accessibility data as previously described in Cusanovich et al [23]. Briefly, a suspension of islet cells were obtained and pelleted 5 min at 4°C 500 × g. The media was aspirated and the cells were washed once in 1 ml PBS. The cells were pelleted again for 5 min at 4 °C 500 × g and then resuspended in 1 ml of cold lysis buffer (10 mM Tris-HCl, pH 7.4, 10 mM NaCl, 3 mM MgCl_2_, 0.1% IGEPAL CA-630, supplemented with 1X protease inhibitors (Sigma P8340)). Nuclei were maintained on ice whenever possible after this point. 10 μl of 300 μM DAPI stain was added to 1 ml of lysed nuclei for sorting. To prepare for sorting, 19 μl of Freezing Buffer (50 mM Tris at pH 8.0, 25% glycerol, 5 mM MgOAc_2_, 0.1 mM EDTA, supplemented with 5 mM DTT and 1X protease inhibitors (Sigma P8340)) was aliquot into each well of a 96-well Lo-Bind plate. 2,500 DAPI^+^ nuclei (single cell sensitivity) were sorted into each well of the plate containing Freezing Buffer. The plate was sealed with a foil plate sealer and then snap frozen by placing in liquid nitrogen. The frozen plate was then transferred directly to a −80 °C freezer. Subsequently, the sample was shipped from NIH to UW overnight on dry ice. The plate was then thawed on ice and supplemented with 19 μl of Illumina TD buffer and 1 μl of custom indexed Tn5 (each well received a different Tn5 barcode). The nuclei were tagmented by incubating at 55 °C for 30 min. The reaction was then quenched in 20 mM EDTA and 1 mM spermidine for 15 min at 37 °C. The nuclei were then pooled and stained with DAPI again. 25 DAPI^+^ nuclei were then sorted into each well of a 96-well Lo-bind plate containing 11.5 μl Qiagen EB buffer, 800 μg/μl BSA, and 0.04% SDS. 2.5 μl of 10 μM P7 primers were added to each sample and the plate was incubated at 55 °C for 15 min. 7.5 μl of NPM was then added to each well. Finally, 2.5 μl of 10 μM P5 primers were added to each well and the samples were PCR amplified with following cycles: 72 °C 3min, 98 °C 30s, then 20 cycles of 98 °C for 10 s, 63 °C for 30 s, 72 °C for 1 min. The exact number of cycles was determined by first doing a test run on 8 samples on a real-time cycler with SYBR green (0.5X final concentration). PCR products were then pooled and cleaned on Zymo Clean&Concentrator-5 columns (the plate was split across 4 columns) eluting in 25 μl Qiagen EB buffer and then all 4 fractions were combined and cleaned using a 1X Ampure bead cleanup before eluting in 25 μl Qiagen EB buffer again. The molar concentration of the library was then quantified on a Bioanalyzer 7500 chip (including only fragments in the 200-1000 bp range) and sequenced on an Illumina NextSeq at 1.5 pM concentration.

#### QC and pre-processing

##### Step 1. Barcode correction and filtering

Each barcode consists of four 8-bp long indexes (i5, i7, p5, and p7). Reads with barcode combinations containing more than 3 edit distance for any index were removed. If a barcode was within 3 edits of an expected barcode and the next best matching barcode was at least 2 edits further away, we corrected the barcode to its best match. Otherwise, the barcode was classified as ambiguous or unknown.

##### Step 2. Adapter trimming and alignment

Adapters were removed using Trimmomatic [31] with NexteraPE adapters as input (ILLUMINACLIP:NexteraPE.fa:2:30:10:1:true TRAILING:3 SLIDINGWINDOW:4:10 MINLEN:20) and aligned to hg19 reference using BWA-MEM (v0.7.15-r1140, options: -I 200,200,5000) [25]. The final alignment was filtered using samtools to remove unmapped reads and reads mapping with quality < 10 (-f3 -F3340 -q10) as well as reads that were associated with ambiguous or unknown barcodes.

##### Step 3. Deduplication and nuclei detection

Duplicates from the pruned file were removed using a custom Python script on a per-nucleus basis. Using the distribution of reads per barcode, we applied bi-culstering, as implemented in the mclust [32] R package, to differentiate between background barcodes and barcodes that correspond to a nucleus. Using the list of non-background barcodes, we split the aggregate bam file into constituent bam files corresponding to each barcode representing a single nucleus using a custom Python script.

##### Step 4. Quality assessment of each single nucleus

For each single nucleus, we computed ATAC-seq quality metrics such as fragment length distribution, transcription start site (TSS) enrichment, short-to-mononucleosomal reads ratio, total autosomal reads, and fraction of reads overlapping peaks. We removed nuclei with a) total reads outside 5% to 95% range [34578, 226755] of all the nuclei, and b) TSS enrichment of <2.7 (5%-tile) from further downstream analysis.

##### Step 5. Aggregate sci-ATAC-seq peaks

We pooled reads from filtered barcodes from the previous steps to create an aggregate bam file. Peaks were called and filtered as described previously in the Bulk Islet ATAC-seq analysis section.

### 2.3. Cluster analysis

#### Feature selection and clustering

We generated a list of TSS distal peaks (>5 kb away from the nearest TSS based on RefSeq genes [33]) from the aggregate sci-ATAC-seq data. For each nucleus, we counted the number of reads overlapping the peaks using the Rsubread package [34]. We then adopted a logistic regression approach to remove peaks where binarized accessibility across nuclei was significantly associated (Bonferroni corrected p-value < 0.05) with sequencing depth. This approach should help to reduce the bias associated with sequencing depth, as the remaining peaks are no longer associated with this technical factor, a strategy that has been successfully implemented in single cell RNA-seq data analysis [35]. The resulting count matrix was RPKM-normalized and reweighted using the term-frequency and inverse-document-frequency (TF-IDF) method [13]. To do this, we first weighted all the sites for individual nuclei by the total number of sites accessible in that cell (“term frequency”). We then multiplied these weighted values by log(1 + the inverse frequency of each site across all cells), the “inverse document frequency.” The TF-IDF transformed matrix was then reduced to 30 principal components using Principal Component Analysis (PCA) and used as input to generate a two-dimensional embedding using the Uniform Manifold Approximation Method (UMAP, n_neighbors = 20) [36]. We identified clusters in the two-dimensional embedding in an unsupervised manner using a density-based clustering method (hdbscan, minPts = 20) as implemented in dbscan R package [37].

#### Cell identity assignment

The cell identities were assigned based on de-facto cell-type-specific hormone markers: *INS-IGF2* (beta), *GCG* (alpha), *SST* (delta) etc. A marker gene was said to be present in a nuclei if a read mapped within 5 kb of the GENCODE (v19) gene body annotation [38]. For additional verification of cell-identity, we computed RPKM normalized aggregate ATAC-seq signal across cell-type marker genes reported in an islet scRNA-seq study [17]. Finally, we evaluated the enrichment of cells from each cell-type cluster relative to their expected population proportion using two-sided binomial test across ten bins of sequencing depth (~145 cells/bin).

### 2.4. Deep learning signal and peak upscaling

#### Model

The U-Net model [39] takes input sequences and outputs prediction sequences. The goal of model training is to reduce the error between the prediction output and a representation of ground truth. For signal upscaling, the input sequence is base-wise scores of BAM pileups (read-depth) corresponding to a subsample of n cells (randomly sampled from 600 cells) and the output sequence is base-wise scores of BAM pileups using reads from all 600 cells. Peak upscaling not only uses the subsampled BAM pileup scores as inputs, but it also uses the binary base-wise values from calling peaks with MACS2 on the subsampled BAM alignments. Output sequences for peak upscaling are the binary base-wise values from calling peaks with MACS2 on the data. We created two models, each separately on data from alpha and beta cells. Because both had different number of constituent single-nuclei, we matched the size of output dataset by randomly sampling 600 cells from each cell-type cluster. The input datasets were created by sampling n cells from the set of 600 cells such that the total number of reads is similar across both models.

The network architecture of the U-Net used in this study is illustrated in **Fig. S2A**. It consists of a contracting, convolutional path (left side) and an expansive, deconvolutional path (right side). The contracting path consists of repeated applications of two kernel size 11 convolutions (unpadded convolutions) with rectified linear unit (ReLU) activation, and a kernel size 2 max pooling operation with stride 2 for downsampling. Each downsampling step halves the length of the activation sequence while doubling the number of feature channels. Every step in the expansive path consists of a kernel size 2 deconvolution layer with a linear activation function that halves the number of feature channels, a concatenation with the correspondingly cropped feature map from the contracting path, and two kernel size 11 convolution layers with ReLU activations. The cropping is necessary due to the loss of border sequence steps in non-padded convolution. At the final layer a kernel size 1 convolution with either an ReLU (for signal upscaling) or sigmoid (for peak upscaling) activation function generates the sequence of predictions. Due to the use of unpadded convolutions, the prediction sequence is shorter than the input sequence by a constant border width. Although the U-Net can accept arbitrary length input sequences, we fix all training samples to be of length 6700, which results in output prediction sequences of length 4820. In total, the network has five steps each in the contracting and expansive paths for a total of 27 convolutional layers and 8,998,529 training parameters. The model was implemented using Keras [40] with the Tensorflow [41] backend, and experiments were run using Titan Xp and GTX 1080 Ti GPUs.

To reduce overfitting, we split chromosomes into training, validation, and testing sets. The model was fit using the ADAM optimizer [42] with a learning rate of 1e-5 and a batch size of 128 for 50 epochs. Separate loss functions, and hence models, were used to solve signal and peak upscaling. For signal upscaling, we used the mean squared error base-wise loss function. For peak upscaling, the loss function was the sum of the cross-entropy base-wise loss and the Dice-coefficient loss, also known as F1 score. We used mean average precision, a common evaluator for object detection, and Pearson correlation as the output evaluation metrics for peak and signal upscaling, respectively. This downscaling and model training were repeated for n=5, 10, 28, 50, 100, 200, 300, 400, and 500 cells.

#### Generating upscaled peaks

In order to select a subset of high-confidence peaks from the predicted model output, we adopted a post-hoc approach where we compared the number of cell-type-specific peaks for alpha, beta, and delta cells, and chose a threshold where they had a similar number. For predicted delta cell peaks, we combined the results from alpha and beta models at the same threshold using bedtools [43] intersect (v.2.27.1) after filtering for the chosen threshold.

### 2.5. Cell-type-specific peaks analysis

#### Cell-type-specific peaks

Peaks specific to each cell-type were obtained by comparing peaks in one cell-type with all other cell-types using bedtools.

#### T2D GWAS SNPs enrichment

Enrichment of T2D associated GWAS SNPs from DIAMANTE [3] was tested using GREGOR (v1.3.1) [44]. Specifically, we used the following parameters: r2 threshold (for inclusion of SNPs in LD with the diabetes associated GWAS SNPs) = 0.80, LD window size = 1 Mb, and minimum neighbor number = 500. P-values were adjusted according to Bonferroni threshold for multiple testing burden.

#### fGWAS analysis

We used fGWAS [45] to model shared properties of loci affecting a trait. We ran fGWAS (v0.3.6) with DIAMANTE T2D GWAS summary data and cell-type ATAC-seq peaks from three cell types as input annotations. For each individual annotation, the output model provided maximum likelihood enrichment parameters and annotations were considered as significantly enriched if the parameter estimates and 95% confidence interval (CI) did not overlap zero. We then used fGWAS to run a conditional analysis in a pair-wise manner where enrichment of one model was evaluated conditional on the output models from other annotations.

### 2.6 Cicero co-accessibility analysis

In order to link TSS distal ATAC-seq peaks with target genes, we used Cicero [46], which identifies co-accessible pairs of DNA elements using single-cell chromatin accessibility data. We used these results to infer connections between regulatory elements and their target genes. We ran Cicero (v1.0.15, default parameters) with cells from the alpha and beta cell clusters separately. To do this, we first called peaks on each cluster and counted the number of reads per nuclei overlapping the peaks. The resulting count matrix was used as input to Cicero along with the UMAP projection for each cluster. Finally, in order to decide a threshold for filtering co-accessible peak pairs, we computed Fisher odds ratio for enrichment of co-accessible peaks versus distance matched non co-accessible peaks (co-accessibility < 0) with three different three-dimensional chromatin looping data sets: islet Hi-C [47], islet promoter capture Hi-C (pcHi-C) [48], and EndoC Pol2 ChIA-PET anchors [49]. For overlap, we checked if both the ends of the Cicero loops intersected with both the anchors from the experimental chromatin looping data. Public epigenome browser session links have been included in **Table S7**.

#### T2D GWAS SNP overlap analysis

In order to link T2D GWAS SNPs with the target genes, we utilized 380 independent GWAS signals from DIAMANTE that were genetically fine-mapped to 99% credible set. We filtered SNPs within each set to have >0.05 posterior probability of association (PPAg). We then checked for each GWAS signal whether SNPs passing the criteria mapped within 1 kb of cell-type-specific ATAC-seq peaks. To obtain Cicero target genes, we checked if an ATAC-seq peak was a) within 1 kb of a variant, b) outside 1 kb range of a RefSeq TSS, and (c) linked to an ATAC-seq peaks that was within 1 kb of a RefSeq TSS. The binary overlap matrix was clustered using hierarchical clustering with binary distance method.

## 3. RESULTS

### 3.1. sci-ATAC-seq captures tissue relevant characteristics similar to bulk ATAC-seq

Pancreatic islets represent approximately 1-2% (by mass) of total pancreatic tissue [1] and therefore requires specialized approaches to isolate in a manner that maintains viability. We obtained a highly pure (>95% purity and >92% viability) sample of human pancreatic islet tissue from one individual (cadaveric donor, female, 43 years old, and non-diabetic) and profiled chromatin-accessibility using sci-ATAC-seq protocol [23] as described previously (**Fig. 1A**, **Table S2**). In total, we obtained 1,690 single-cell ATAC-seq datasets with depth ranging from 17,667 to 415,237 (median: 79,482) reads per nucleus, and TSS enrichment from 0.77 to 9.80 (median: 3.91) after removing background barcodes (**Fig. S1A**). For quality assessment of each single nucleus, we reasoned that total reads and TSS enrichment values are more suitable metrics for identifying nuclei with poor signal-to-noise ratio than using fraction of reads in peaks as the latter may bias counts for under-represented cell-type populations in the analysis (**Fig. S1B-C**). Based on these criteria, we obtained high-quality sci-ATAC-seq data for 1,456 single-nuclei. In addition to sci-ATAC-seq data, we generated high-quality bulk ATAC-seq data for ten islet samples with >47 M reads and >4.4 TSS enrichment per sample (**Table S3)**. Using our approach to identify high-confidence (master) peak calls across samples (see methods), we obtained 106,460 bulk islet accessible chromatin peaks.

**Figure 1.**
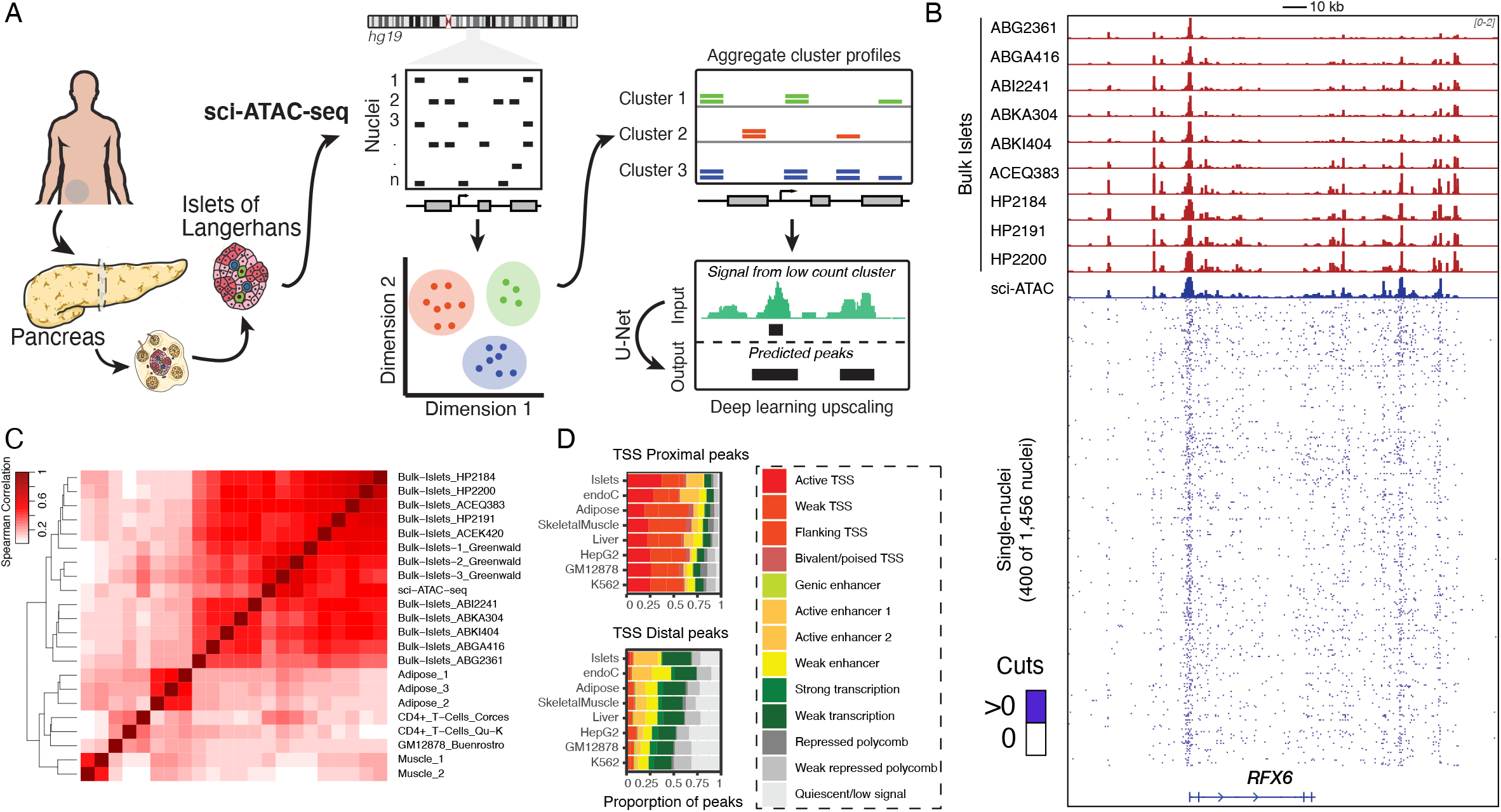
Schematic of sci-ATAC-seq study. (A) sci-ATAC-seq protocol for generating single-nuclei ATAC-seq data from a pancreatic islet sample. The data is then used to identify constituent cell-types and use deep-learning model to predict peaks on the clusters with fewer nuclei count. (B) ATAC-seq signal tracks for 10 bulk islet samples and sci-ATAC-seq islet sample. Bottom tracks show the signal across a random subset of upto 400 single-nuclei. Signal tracks are normalized to one million reads and scaled between 0-2. (C) Spearman correlation between aggregate sci-ATAC-seq, 13 bulk islets, 3 adipose, 2 muscle, 2 CD4+ T-cells, and 1 GM12878 sample (see **Table S1**). (D) Distribution of aggregate sci-ATAC-seq TSS proximal and distal peaks across bulk islet derived ChromHMM segmentations.

We then compared the aggregate islet sci-ATAC-seq data with bulk ATAC-seq samples from islets and other tissues. For this, we called 156,311 peaks on the aggregate sci-ATAC-seq. We found that aggregate sci-ATAC-seq profiles were concordant and clustered together with the other bulk islet samples indicating that aggregate sci-ATAC-seq can capture chromatin accessibility in a manner equivalent to bulk ATAC-seq assays (**Fig. 1B-C**, **S1D-E**). Further, to understand if the aggregate sci-ATAC-seq peaks capture islet-specific regulatory features, we compared the distribution of peaks across chromHMM chromatin state maps in eight tissues, including islets and the EndoC human beta cell line [8]. We found that islet sci-ATAC-seq peaks overlap active TSS and active enhancer segmentations in islet and EndoC (a beta cell line) chromatin state maps to a larger extent compared to other tissues (**Fig. 1D**). Because chromHMM enhancer states are driven by H3K27ac marks and are known to be associated with tissue-specific enhancer activity [5,50], our results indicate that sci-ATAC-seq data capture the underlying islet-specific chromatin architecture similarly to bulk islet ATAC-seq assays. Overall, these results indicate that our aggregate islet sci-ATAC-seq data are of high quality and suggests that the underlying individual nuclei could reveal valuable cell-specific patterns of the constituent cell-types.

### 3.2. sci-ATAC-seq reveals constituent cell-types in pancreatic islets

The aggregate sci-ATAC-seq profile of the islet is constituted of signal from distinct cell-types. For identifying these cell-types, we leveraged the observation that TSS distal regions capture cell-type-specific accessibility patterns and are effective at classifying constituent cell-types [51]. We adopted a multi-step process to robustly detect and identify islet subpopulations (see methods, **Table S4**). This approach produced four distinct clusters (**Fig. 2A**). In order to assign a cell-type identity to the clusters, we merged nuclei in each cluster to create aggregate chromatin accessibility profiles and systematically examined the patterns of accessibility at multiple cell-type marker loci. We found three clusters to have distinct chromatin-accessibility patterns at *GCG, INS-IGF2*, and *SST* loci corresponding to three major islet cell-types: alpha, beta, and delta cells (**Fig. 2B**). The fourth cluster (95 nuclei, ~7% of all nuclei) showed a “mixed” cell-type appearance as shown by signal at multiple cell-specific markers. We reasoned that these are likely to be nuclei doublets resulting from barcode collisions inherent to the combinatorial indexing protocol, and thus should have skewed ATAC-seq read coverage. Indeed, we observed that nuclei assigned to the mixed cell cluster were significantly (nominal P-value = 7.3e-7, binomial test) enriched in the high sequencing depth bin relative to nuclei from other clusters (**Fig. 2C**). As such, these nuclei were removed from further analyses yielding a total of 1,361 nuclei with 51%, 47%, and 2% assigned to beta, alpha, and delta cell-type respectively. These estimates agree with the existing estimates of pancreatic islet cell-type proportions observed in confocal microscopy or single-cell transcriptomics experiments [17,52–54]. As additional validation of our cell-type assignments, we used cell-type signature genes from a published islet scRNA-seq study [17] and observed cluster-specific chromatin accessibility consistent with our assigned cell identities (**Fig. 2D-E**).

**Figure 2.**
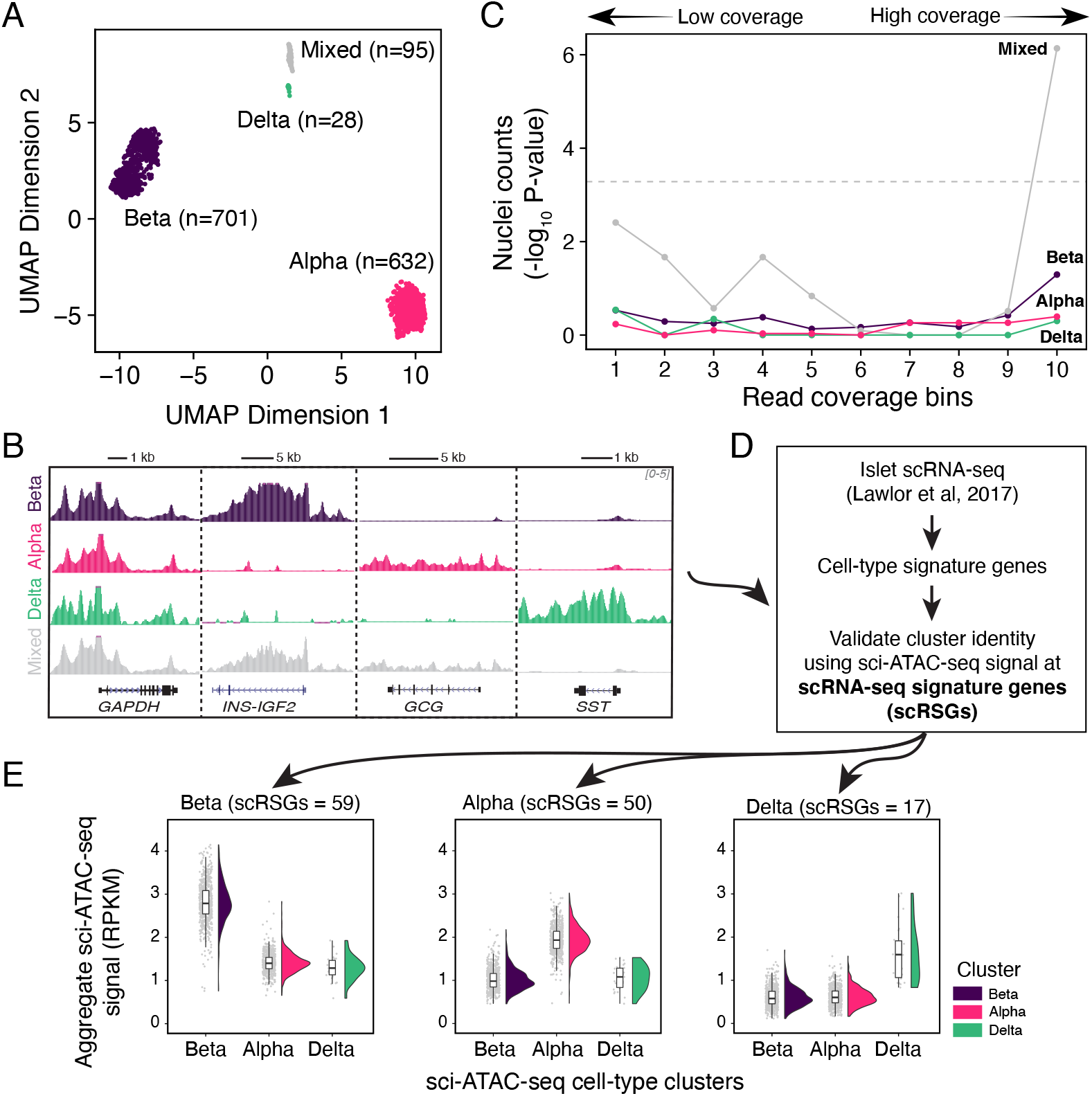
Clustering and identification of cell-type clusters in sci-ATAC-seq data. (A) UMAP projection with clustering of 1,456 single-nuclei islets represented by each single point into four clusters as identified by density-based clustering. (B) Enrichment of cells from each cluster relative to their expected population proportion across different read sequencing depth bins. Sequencing depth increases with the bin number. (C) Genome browser tracks showing signal at different cell-type marker loci: alpha (*GCG*), beta (*INS-IGF2*), delta (*SST*), and a housekeeping gene *(GAPDH)*. Tracks are normalized to one million reads and scaled between 0-5. (D) Overview of independent cluster verification scheme utilizing cell-type signature genes as identified by an islet scRNA-seq study by Lawlor et al (2017). (E) Plot of aggregate ATAC-seq signal (normalized using RPKM) at scRNA-seq derived cell-type signature genes for alpha, beta, and delta cells. Number of signature genes for each cell type indicated in the legend.

We then analyzed the chromatin accessibility profile for each cell-type cluster. For this, we aggregated nuclei within each cluster and identified peaks using MACS2. We identified 129,046 sites for alpha and 120,116 sites for beta cells. However, because the delta cluster had only 28 cells (corresponding to ~2 M reads), we reasoned that MACS2 would not perform ideally on data with such low depth. Indeed, we only identified 49,293 peaks using MACS2 on the delta cell aggregate reads.

### 3.3. Deep learning enables robust peak calls on less abundant delta cells

To solve the challenge of learning cell-type-specific features from the sparse signal in low count delta cell cluster, we used a deep learning approach based on U-Net architecture (**Fig. S2A**). U-Net was first developed for biomedical image segmentation but has since been applied to many other problems including audio and image super resolution. Its use in the super resolution problem served as the main impetus for our choice of model to upscale genomic signals. We formulated our approach as a classification problem where we use sparse signal and corresponding peak calls (equivalent to a low-resolution image) to predict dense and high-quality peak calls (equivalent to a high-resolution image). In order to avoid overfitting, we adopted a rigorous training scheme. We divided the chromosomes into training, validation, and testing sets (**Fig. 3A**, **S2B**), and tested the performance of models within the same cell-type and across different cell-types. We reasoned that our islet sci-ATAC-seq data is an ideal fit for this problem as all the nuclei come from the same individual and processing batch and should, therefore, contain no genetic or technical biases that would influence within or across cell-type predictions. Since we had high-quality data from two cell-types, we trained two models: one model was trained using 28-cell and 600-cell data from alpha cells (alpha trained model), while the second model was trained similarly on data from beta cells (beta trained model). We then compared peak predictions from both models to corresponding MACS2 peaks from the 600-cell data. We found that results from cross cell-type predictions of both models outperformed MACS2 peak results as measured by mean average precision **(Fig. 3B)**, suggesting that the U-Net was able to reconstruct peak calls from sparse signal independent of the specific cell-type it was trained on. We highlight several examples where the model was able to successfully predict peaks that were absent in sparse 28-cell data but present in 600-cell data of a cell-type **(Fig. 3C)**. These predictions could not have transferred or “copied over” from the training data because the training cell-type had no signal or peak call at the given locus. Based on these results, we decided to use the U-Net models to predict peaks for the low-count delta cell cluster. Since the U-net model provides a posterior probability score for each peak call prediction, we sought to create a high-confidence set of predicted peak calls for each cell-type. We used a threshold of 0.625 to filter predicted peaks for each cell-type. The choice of threshold was used to control for potential false positives and final number of predicted cell-type peaks (**Fig S2C-D**).

**Figure 3.**
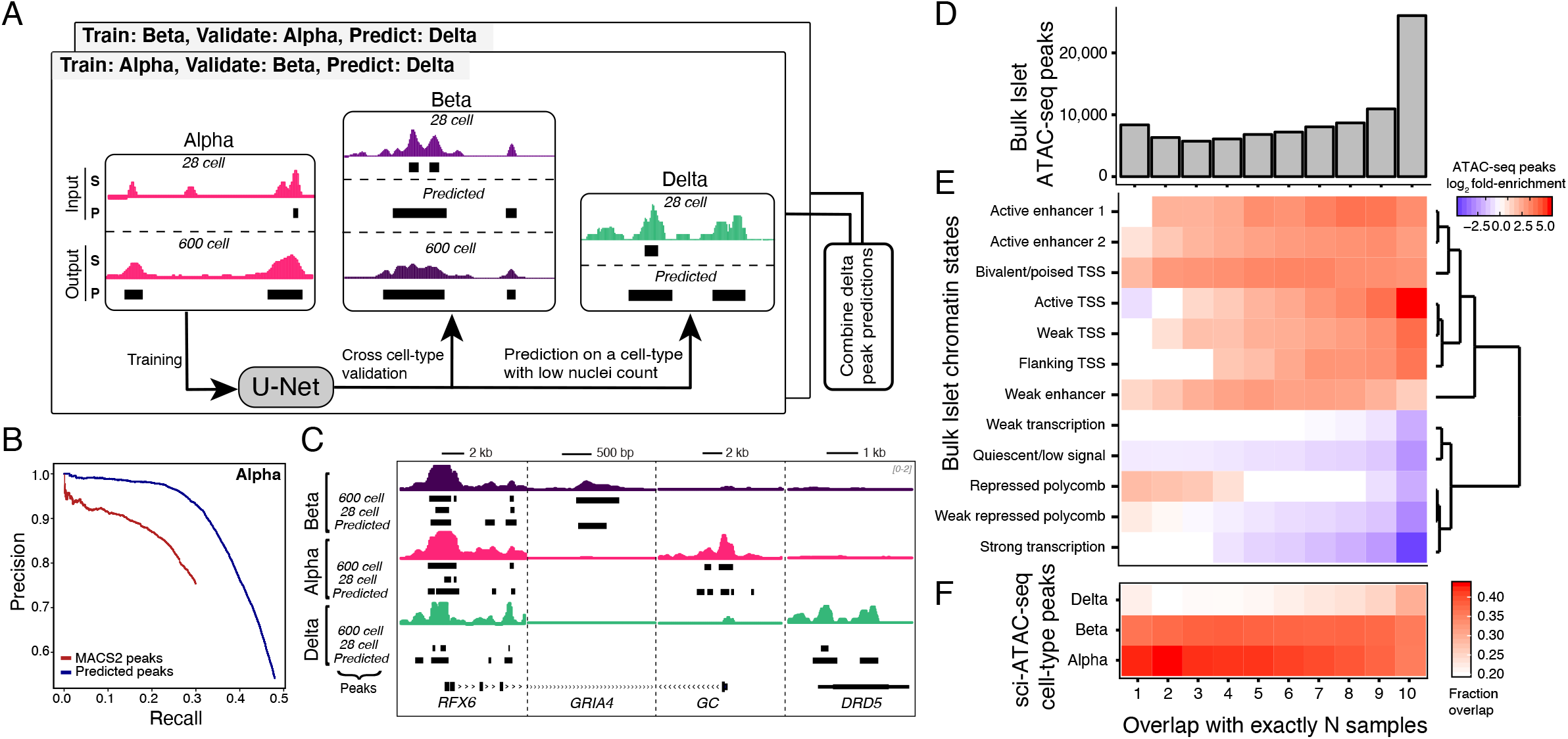
Deep learning upscaling from sparse low-count nuclei clusters. (A) Schematic of U-Net training scheme. Two models are depicted in the illustration: one trained on alpha cells data as input and other trained on beta cells as input. Delta cell peak predictions from both models are combined to get final predictions (see Methods). (B) Precision-recall curve comparing peak calls from MACS2 on downscaled data (alpha cell-type) with predicted peak calls from 28 cell U-net model (trained on beta, predicted on alpha). (C) Example loci illustrating peak upscaling with the model. For each cell-type, four tracks are shown: full signal track, peak calls on full data, peak calls on subsampled data, and predicted peak calls. The predicted peak calls are obtained from a model trained on a different cell-type. For delta predicted peak calls, intersection of prediction from both alpha and beta models are shown. Signal tracks normalized to one million reads and scaled between 0-2. (D) Reproducibility of master peaks from bulk islet ATAC-seq across individual samples. (E) Fold enrichment (log_2_) of different sets of reproducible peaks from bulk islet ATAC-seq across 13 islet chromatin states. Genic Enhancer is not shown because of no enrichment. (F) Overlap of cell-type peaks (alpha, beta, predicted delta) with different sets of reproducible peaks from bulk islet ATAC-seq data.

Further, considering that delta peak predictions from both alpha and beta model were highly concordant (Jaccard index of 0.85), we used the intersection of the results as the final predicted outcome. Now with representative peak calls from each cluster, we compared them to bulk islet ATAC-seq master peak calls (**Table S5**). We found that master peaks derived from bulk islets were highly reproducible across samples, with >70% of peaks occurring in five or more of the ten samples (**Fig. 3D**). Predictably, we also observed that the chromatin states corresponding to “Active TSS” and “Active Enhancer” showed enrichment with increasing reproducibility of master peaks. Likewise, chromatin states such as “Repressed polycomb”, “Weak transcription”, “Quiescent/low signal” showed a depletion with increasing islet ATAC-seq peak reproducibility (**Fig. 3E**). Similarly, when we compared cell-type peaks to the master peaks, we found that the proportion of peaks from each cell-type increased with the increasing reproducibility of bulk peaks, suggesting that highly reproducible peaks are driven by all constituent cell-types while peaks that occur in fewer samples might originate from underlying cell-population variability (**Fig. 3F)**.

While the primary model of our interest was trained using data from 28 cells to predict 600-cell equivalent peaks, we asked if the model would perform similarly for a varying resolution of input data. For this, we subsampled cells from alpha and beta cell clusters to sets of different cell counts, starting with as few as five cells to 500 cells. We found that the performance of the model increased with the increasing number of cells used in the input training data (**Fig. S2E**). There was up to five-fold gain in coverage of T2D GWAS SNPs in beta predicted peaks compared to MACS2 peaks (**Fig. S2F**) even when fewer cells were used as input training data (**Fig. S2G)**. These results suggest that the deep-learning strategy is applicable to a range of input data typically seen in single-cell sequencing experiments.

Overall, our results show that deep-learning driven feature prediction can help us recover tissue and cell-type relevant chromatin accessibility patterns from sparse and noisy data. Using this approach can enhance biological discoveries, which is challenging with rare cell populations.

### 3.4. T2D GWAS enrichment at cell-type-specific chromatin signatures

We computed the overlap enrichment of T2D GWAS loci in cell-type peak annotations from alpha, beta, and delta cells using a Bayesian hierarchical model, as implemented in fGWAS [45]. fGWAS allows calculation of marginal enrichment associations for one cell type conditioned on another by using not only the subset of genome-wide significant loci but also the full genome-wide association summary statistics. We observed that annotations from all three cell types were highly enriched for T2D GWAS loci, with beta-cell annotations having the highest enrichment values. (**Fig. 4A**). However, when we accounted for marginal associations using a joint model, we found that beta cells are the only cell type to remain enriched after adjusting for the other two cell types. This result suggests that shared or beta cell-specific chromatin accessibility peaks drive the association with T2D GWAS. More broadly, these findings illustrate how single cell chromatin profiling results, when coupled with conditional statistical enrichment analyses, can dissect specific cell types that drive enrichment in bulk tissue samples.

**Figure 4.**
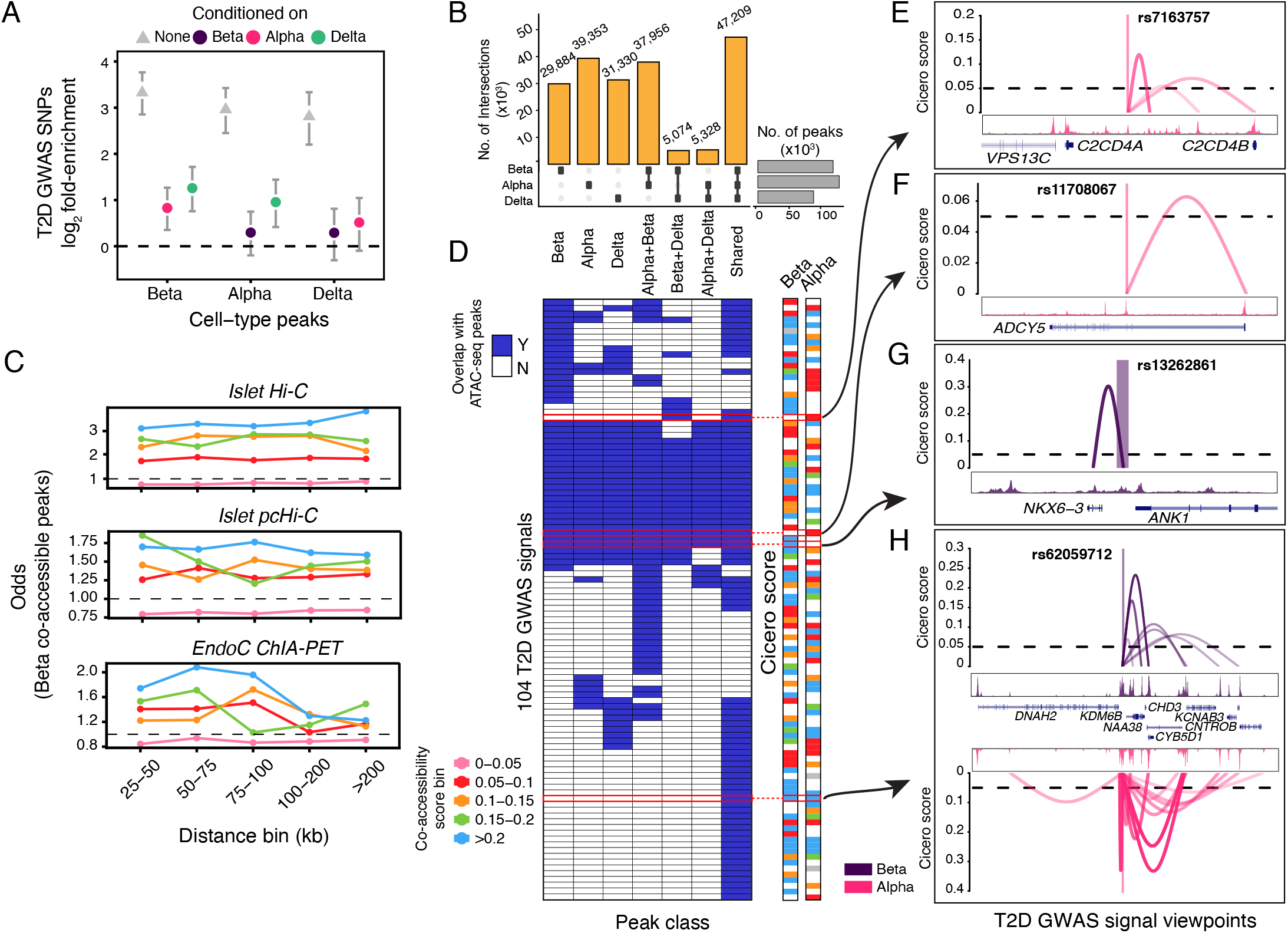
Enrichment of T2D GWAS signals in cell-type-specific chromatin and linking them to target genes. (A) Fold enrichment (log_2_) of T2D GWAS SNPs in cell-type peaks in single and conditional analysis mode using fGWAS tool. For each cell-type, three enrichment values with 95% confidence intervals are shown: None (single annotation mode), Alpha (conditioned on alpha), Beta (conditioned on beta), and Delta (conditioned on delta). (B) Partitioning of alpha, beta, and predicted delta peaks in mutually exclusive sets of cell-type-specific peaks. The subplot (on right) shows the total number of peaks for each cell-type. (C) Distance-matched Fisher odds that beta cell co-accessibility links overlap islet Hi-C, islet pcHi-C, and ChIA-PET chromatin loops across different co-accessibility threshold bins. (D) Overlap of T2D GWAS credible set SNPs with cell-type-specific peaks. Bin is colored if there’s at least one SNP (PPAg > 0.05) in the 99% genetic credible set of the T2D GWAS signal located within 1 kb of an ATAC-seq peak. Cicero score columns are colored to indicate the score of the highest scoring link to the target gene. (E) Viewpoint plot of alpha Cicero connections centered at rs7163757 for *C2CD4A/B* locus, (F) alpha Cicero connections centered at rs11708067 for *ADCY5* locus, (G) beta Cicero connections centered at rs13262861 for *ANK1* locus, and (H) Cicero connections for both alpha and beta centered at rs62059712 for *ATP1B2* locus. The viewpoint region is +/− 1kb of the region from the variant.

We next partitioned the peaks into exclusive sets based on the cell-types shared by each peak. Because the delta cell cluster has fewer reads compared to alpha and beta cells, we did not utilize read count-based approaches to determine cell-type-specific peaks. Instead, we used peak level metrics to identify peaks exclusive to a combination of cell types. We found that a majority of peaks (47,209) were shared across all cell-types and that each cell type had a set of unique accessible sites (29,884 beta, 39,353 alpha, 31,330 delta) (**Fig. 4B**). Consistent with our expectations, TSS-proximal shared peaks mostly overlapped active TSS chromatin states compared to cell-type-specific peaks which had a larger proportion of peaks in active enhancer states (**Fig. S3A)**. We then used a complementary enrichment approach with the GREGOR tool [44] to determine if T2D GWAS loci are enriched in each subclass of peaks. We found that T2D GWAS loci were highly enriched in shared peaks (P-value 1.64e-16, fold enrichment 2.03) and beta cell-specific peaks (P-value 6.42e-6, fold enrichment 1.91) (**Fig. S3B**). We also observed moderate enrichment of T2D GWAS SNPs in other sets of cell-type-specific peaks, but strikingly, there was little enrichment in delta cell-specific peaks (P-value 3.12e-3, fold enrichment 1.55) and no significant enrichment in alpha cell-specific peaks (P-value 1.83e-01, fold enrichment 1.16). This suggests that the role of alpha and delta cells in the mechanisms underlying genetic predisposition to T2D pathophysiology might be limited compared to beta cells. Further, the enrichment of shared peaks also suggests that these variants might be affecting regulatory pathways that are shared across all cell types in the islets, suggesting that a narrow focus on studying beta cells only for T2D might not yield a complete picture of the disease mechanisms. These independent enrichment findings from the GREGOR tool are consistent with the results from the fGWAS analysis (**Fig. 4A**), indicating the robust nature of these results.

### 3.5. Linking cell-type-specific chromatin accessibility to target genes

One of the primary challenges in understanding the underlying biological mechanisms at non-coding T2D GWAS variants is the identification of their target genes. Risk variants occurring in enhancer regions can often interact with their target genes that are not adjacent. Multiple studies have examined the regulatory landscape of pancreatic islets and relevant cell lines using chromosome conformation capture techniques to nominate target genes [47–49]. However, most of these studies were conducted on bulk islet samples, thereby obscuring any cell-specific signatures of chromatin looping. Additionally, chromatin looping studies tend to have noisy signals when two regions are close in linear space, which leads to a bias towards detecting longer-range interactions. In order to mitigate these limitations, we adopted a recently published approach, Cicero [46], which leverages profiles of chromatin co-accessibility across single cells to infer pairs of chromatin peaks that are likely to be in close physical proximity. For this analysis, we focused on alpha and beta cell-types as they were the clusters with the most nuclei. In order to filter the Cicero co-accessible scores for those peak pairs that are more likely to represent true looping, we compared our results to experimentally-defined loops from three independent chromatin looping data sets: islet Hi-C [47], islet promoter capture Hi-C (pcHi-C) [48], and EndoC Pol2 ChIA-PET [49] loops. We found that Cicero peak-pairs from our sci-ATAC-seq data with score >0.05 were strongly enriched to be called as loops in each of the three reference data sets (**Fig. 4C**, **S4A**). With this threshold, we found 190,176 beta cell and 147,716 alpha cell co-accessible peak-pairs.

Using our new catalog of Cicero-inferred chromatin loops, we next sought to link TSS-distal T2D GWAS variants to target gene promoters. We focused on the latest T2D GWAS results and used SNPs in association signals that were genetically fine-mapped to be in a 99% credible set and had >0.05 posterior probability of association (PPAg) [3]. For this mapping procedure, we required that the credible set SNP was not within 1 kb of an annotated TSS and that the other end of the chromatin loop occurs within 1 kb of an annotated TSS. Using this approach across both alpha and beta cells, we found that of the 265 independent GWAS signals containing SNPs that met our criteria, we were able to nominate target genes at 104 of them **(Fig. 4D)**. In a similar manner, we checked if the SNPs within each locus overlapped a cell-type-specific peak (**Table S6**). We observed several notable examples. At the *C2CD4A/B* locus, we found rs7163757 (PPAg 0.095) to be linked to *C2CD4B* in alpha cells (**Fig. 4E).**Using an islet gene expression and genetic integration approach to identify expression quantitative trait loci (eQTL), we previously showed that rs7163757 is associated with *C2CD4B* expression [8], and a subsequent functional study corroborated these findings [55]. At a different locus, we found rs11708067 (PPAg 0.79), located in an islet enhancer within the *ADCY5* gene, to be linked to the TSS of the corresponding gene **(Fig. 4F).** The risk allele of rs11708067 has been reported to be associated with reduced expression of *ADCY5* [56] and functional validation experiments show association with impaired insulin secretion [4,9]. As an example of a beta cell-specific connection, we found variant rs13262861 (PPAg 0.97) within the *ANK1* locus to be linked to nearby *NKX6-3* (**Fig. 4G).** We have previously used islet eQTL data to nominate *NKX6-3* as an islet target gene at this locus [4,8]. The extensive support from previous publications for these three loci serves as positive controls for our results and reinforces the quality of this sci-ATAC-seq data and analyses. Finally, we highlight rs62059712 (PPAg 0.34) within the *ATP1B2* locus as an example of a variant linked to multiple gene promoters across both beta and alpha cell-types **(Fig. 4H)**. Notably, of the 104 T2D GWAS signals for which we were able to nominate target genes in either cell type, 60 (~58%) had more than one nominated target gene.

## 4. DISCUSSION

Single-nuclei chromatin accessibility profiling provides a unique approach for mapping of cell-type-specific regulatory signatures. Here, we utilized the sci-ATAC-seq protocol to generate and study chromatin accessibility profiles for 1,456 high-quality nuclei from a purified pancreatic islet sample. Our dataset and analyses provide high-quality maps of cell-type accessibility profiles and regulatory architecture using an unbiased approach compared to prior maps from sorted cell-type populations. However, it is essential to emphasize that single-cell data present unique challenges, and that our study, which analyzed only one pancreatic islet sample, may be limited in how it can address some of them.

First, de-novo identification of cell types from the sparse single-cell chromatin accessibility data continues to be a challenge. We adopted several strategies to address potential biases in our analysis. Our logistic regression approach to eliminate read depth as a confounding technical variable, combined with the binomial counting strategy to infer doublet enrichment in clusters, enabled us to identify three major cell-type populations corresponding to alpha, beta, and delta cells. In order to assign these cell identities, we relied not only on classical hormone markers, but we also leveraged findings from an independent islet single-cell RNA-seq study to validate our results. While islets have been reported to contain other rarer cell-type populations (<5% of all islet cells) [52], our ability to observe them was limited due to the size of our dataset.

Second, we faced the challenge of identifying reliable cell-specific accessibility patterns across all cell types due to the relatively low abundance of delta cells. As such, our U-Net-based deep learning approach presents a novel strategy for addressing this particular problem. Our model differs from a related deep learning method, Coda [57], by focusing on single cell ATAC-seq as opposed to bulk histone ChIP-seq data and uses a more complex architecture (U-Net) which has been used before in image processing related tasks [39,58] but seen little mention in genomics [59]. We demonstrated, using alpha and beta cells as reciprocal training and testing datasets, that our model successfully learns to predict high-quality peak calls from low cell count data. We observed, however, that there are diminishing returns from using deep learning models when 200 or more cells are used as input to the model, an observation consistent with the threshold of experimental reproducibility highlighted in a recent large-scale single nuclei ATAC-seq study [15]. This consistency with an independent study reinforces the value of our deep learning approach but also highlights a limitation of our delta peak predictions which derive from a low cell count input dataset. Nonetheless, we envision that our method will be useful in scenarios where it is challenging or cost-prohibitive to obtain specific cell populations.

Overall, an important implication of our findings comes from our ability to generate cell-specific chromatin accessibility maps and to infer looping connections from accessible regions to target genes of T2D GWAS variants. A recent T2D GWAS [3] reported >400 independent association signals, but the molecular mechanisms underlying these signals is known only for a subset of the variants. Single nuclei resolution cell-specific regulatory signatures provide a unique opportunity to infer target gene links with non-coding elements. Thus, we integrated cell-type co-accessibility links with T2D GWAS SNPs that were genetically fine-mapped to 99% credible sets to create a higher resolution map of the regulatory landscape underlying 104 distinct T2D GWAS signals. Focusing on the cell-specificity of the chromatin accessibility peaks that anchor these target gene associations, we observed seven classes, representing: i. peaks that are unique to a cell type (three classes), ii. peaks that are shared across all three cell types (one class), iii. peaks that occur in a pair of cell types (three classes). Interestingly, the class of peaks shared across all three cell types comprised 26 of the 104 (25%) T2D GWAS to target gene links even though this class is only one of seven. These results paint a complex picture of disease mechanisms where certain risk variants may mediate target effects through cell-type-specific pathways, while others might affect multiple target genes shared across cell-type populations.

We noted specific examples at the *C2CD4A/B* and *ANK1* loci, where we were able to nominate specific variants linked with islet gene expression and their role in T2D pathophysiology as compelling targets for future mechanistic studies. As this manuscript was under preparation, another similar study appeared as a preprint [60], and as such an important future topic will be to combine and meta-analyze multiple islet single-cell ATAC-seq datasets. Such an endeavor will increase statistical power to detect chromatin features, including loops, at GWAS loci, and eventually enable single-cell resolution chromatin QTL studies, which will help to further narrow in on functional SNPs. Overall, we believe that the data, results, and methodology from this study will be of value to the broader research community.

## DATA AVAILABILITY

The data reported in this paper have been deposited in the dbGaP (accession no. phs001188.v1.p1; FUSION Tissue Biopsy Study—Islet Expression and Regulation by RNAseq and ATACseq). Code for deep learning analysis is available on GitHub at https://github.com/ParkerLab/PillowNet. Custom scripts are available at https://github.com/ParkerLab/islet_sci-ATAC-seq_2019. Additional URLs are provided in **Table S7** of Supplementary Data file.

## ACKNOWLEDGEMENTS

We thank the members of the Parker, Collins, and Shendure labs for helpful suggestions and feedback. This work was supported by American Diabetes Association (ADA) Fellowship to JPD, National Institute of Diabetes and Digestive and Kidney Diseases grant R01 DK117960, National Heart, Lung, and Blood Institute grant U01 HL137182, and the ADA Pathway to Stop Diabetes Grant 1-14-INI-07 to SCJP, and National Institutes of Health Grants 1-ZIA-HG000024 to FSC. The medical art images were obtained from Servier Medical Art by Servier under CC-BY-3.0 license. We thank F. Steemers and L. Christiansen at Illumina for providing indexed Tn5 transposase. We also thank NVIDIA Corporation for donating the Titan Xp GPUs to SCJP.

## AUTHOR CONTRIBUTIONS

FSC, JS, and SCJP conceived the study; MRE, DC, and RMD generated the data; LSZ, JPD, and NN performed initial analysis; VR, DXQ, and SCJP analyzed the data and performed the research presented here; DXQ and YG implemented the U-Net strategy and wrote the code; FSC, JS, and SCJP jointly supervised the work; and VR, DXQ, and SCJP wrote the manuscript with feedback from all the authors.

## SUPPLEMENTARY FIGURE LEGENDS

**Figure S1. ATAC-seq metrics of nuclei from sci-ATAC-seq.** (A) Distribution of reads per barcodes shown with the threshold chosen for filtering background barcodes. (B) Fraction of reads in peaks versus TSS Enrichment, and (C) Total autosomal reads versus TSS enrichment for all single nuclei. Density units are arbitrary. (D) TSS coverage of aggregate sci-ATAC-seq, and (E) Fragment length distribution of aggregate sci-ATAC-seq compared with ten bulk islet ATAC-seq samples.

**Figure S2. Peak calling using deep learning approach.** (A) Schematic of U-Net learning strategy. (B) The training, testing, and validation scheme used for training the models delineating which chromosomes were part of what dataset. (C) Number of predicted peaks (from 28-cell trained model) for each cell type with different output posterior probability thresholds. (D) Number of cell-type specific peaks for alpha, beta, and delta after partitioning into mutually exclusive sets (see methods) with different output posterior probability thresholds. (E) Average precision in predicting peaks compared for all four models (two training and two prediction datasets) with different sizes of input training data. (F) Enrichment of T2D GWAS SNPs (N=378) in predicted beta peak calls (from alpha-trained model) compared with peaks calls from MACS2 on the data with varying size of input training data. (G) Precision and recall curves comparing predicted beta peaks (from alpha-trained model) for varying size of input training data.

**Figure S3.** (A) Distribution of TSS proximal and distal peaks (>5kb from nearest Refseq TSS) in shared peaks and peaks assigned only to alpha, beta, and delta cell-type. (B) Enrichment of T2D GWAS SNPs (N=378) across all cell-type-specific sets of peaks.

**Figure S4.** (A) Fisher odds score for enrichment of alpha co-accessible sites in loop anchors from three different dataset: Islet Hi-C, Islet pcHi-C, and EndoC ChIA-PET.

