## Supplementary Figures for "Single cell ATAC-seq in human pancreatic islets and deep learning upscaling of rare cells reveals cell-specific type 2 diabetes regulatory signatures"

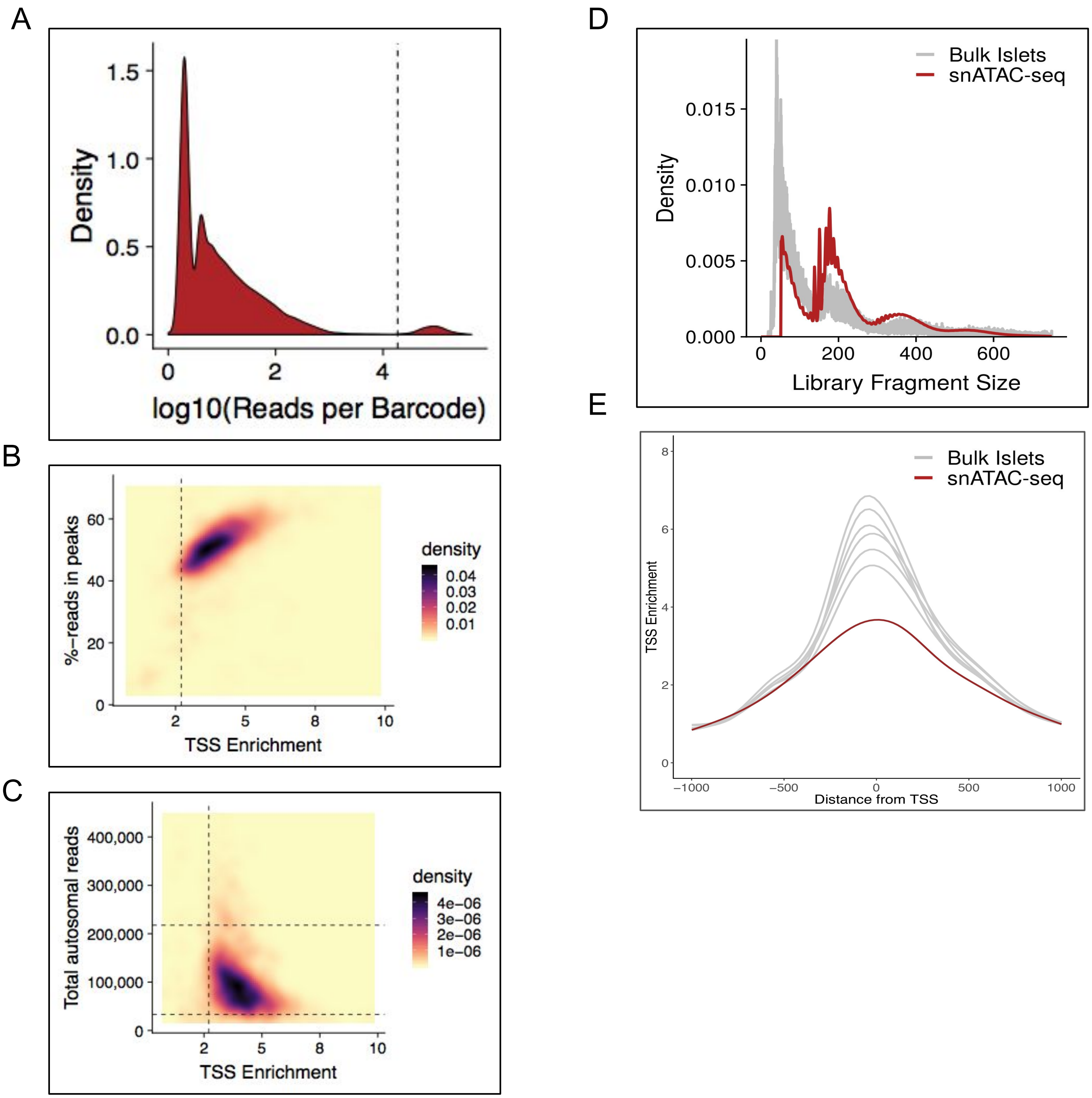


**Figure S1. ATAC-seq metrics of nuclei from sci-ATAC-seq.** (A) Distribution of reads per barcodes shown with the threshold chosen for filtering background barcodes. (B) Fraction of reads in peaks versus TSS Enrichment, and (C) Total autosomal reads versus TSS enrichment for all single-nuclei. Density units are arbitrary. (D) TSS coverage of aggregate sci-ATAC-seq, and (E) Fragment length distribution of aggregate sci-ATAC-seq compared with ten bulk islet ATAC-seq samples.

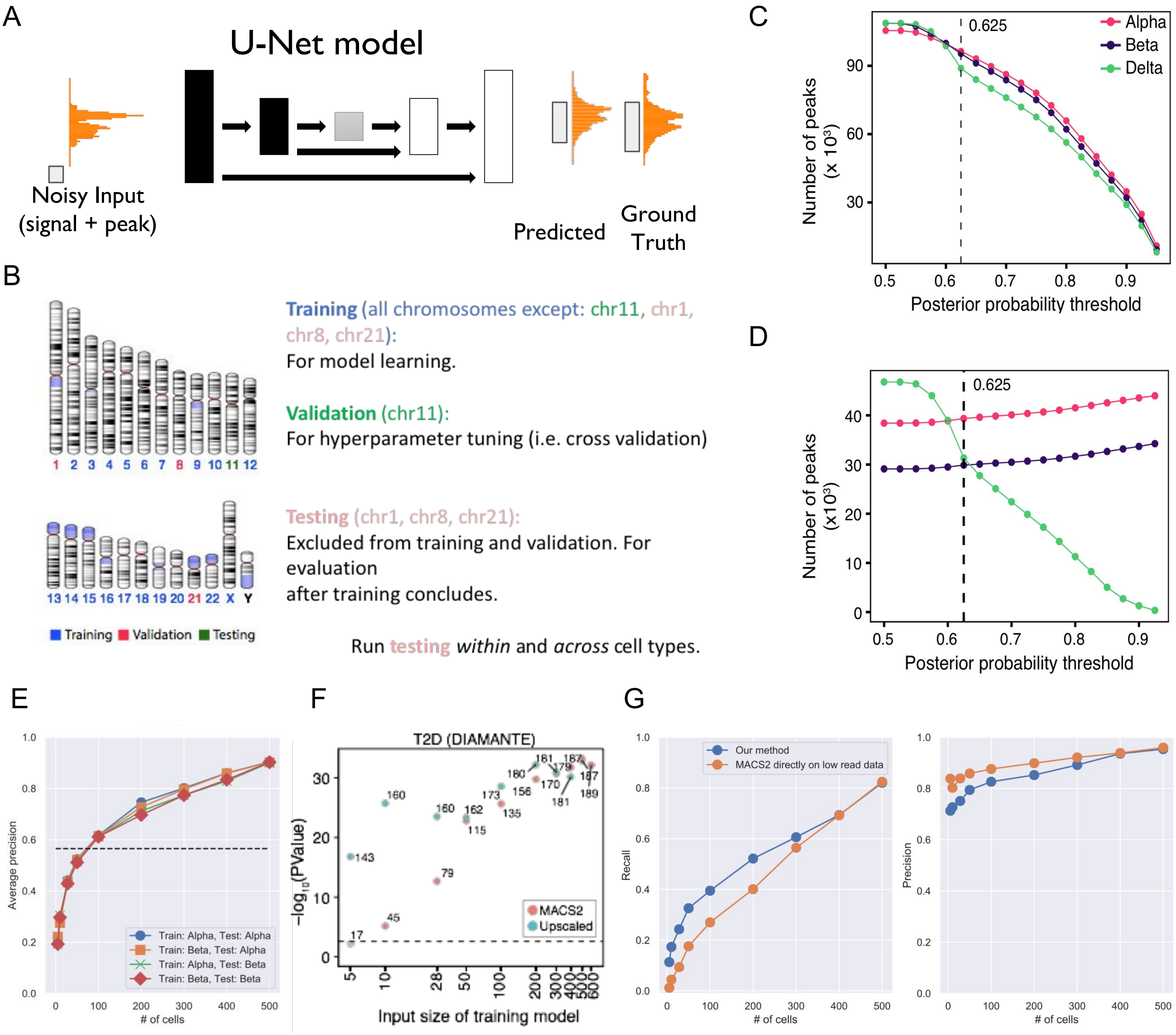


**Figure S2. Peak calling using deep learning approach.** (A) Schematic of U-Net learning strategy. (B) The training, testing, and validation scheme used for training the models delineating which chromosomes were part of what dataset. (C) Number of predicted peaks (from 28-cell trained model) for each cell type with different output posterior probability thresholds. (D) Number of cell-type specific peaks for alpha, beta, and delta after partitioning into mutually exclusive sets (see methods) with different output posterior probability thresholds. (E) Average precision in predicting peaks compared for all four models (two training and two prediction datasets) with different sizes of input training data. (F) Enrichment of T2D GWAS SNPs (N=378) in predicted beta peak calls (from alpha-trained model) compared with peaks calls from MACS2 on the data with varying size of input training data. (G) Precision and recall curves comparing predicted beta peaks (from alpha-trained model) for varying size of input training data.

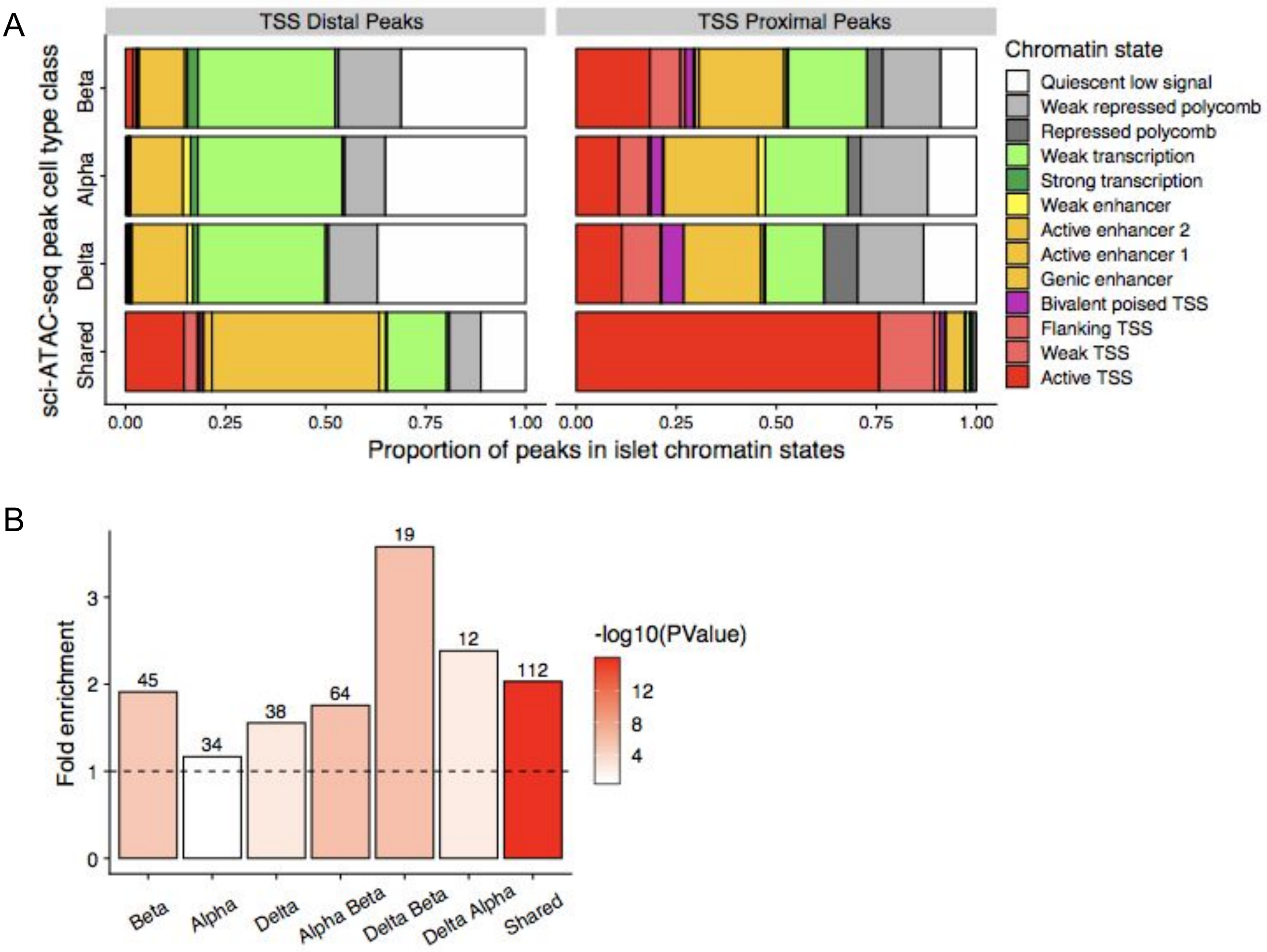


**Figure S3.** (A) Distribution of TSS proximal and distal peaks (>5kb from nearest Refseq TSS) in shared peaks and peaks assigned only to alpha, beta, and delta cell-type. (B) Enrichment of T2D GWAS SNPs (N=378) across all cell-type specific sets of peaks.

A

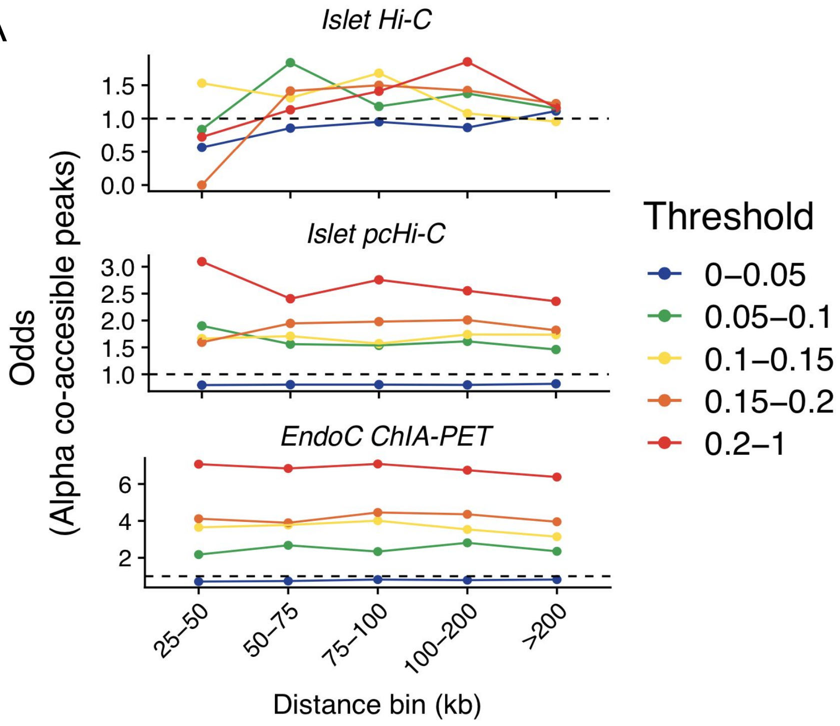


**Figure S4.** (A) Fisher odds score for enrichment of alpha co-accessible sites in loop anchors from three different dataset: Islet Hi-C, Islet pcHi-C, and EndoC ChIA-PET.
